# Neural and behavioural dynamics of the individual and collective self(s)

**DOI:** 10.1101/2023.03.22.533659

**Authors:** David Huepe, Andrés Canales-Johnson, Álvaro Rivera-Rei, Agustín Ibanez, Tristan A. Bekinschtein

**Author notes:** These authors contributed equally to this work.

## Abstract

Usually considered as internal representations of self-concepts, the individual-self and the collective-self have been primarily studied in social and personality psychology while the experimental and theoretical advances of the cognitive and neurophysiological mechanisms of these self-representations are poorly understood. Two competing hypotheses emerge to understand their structure: first, each self-representation corresponds to a specific and independent dimension of self-concept and is therefore conceptualized as separate cognitive components and different brain networks are predicted; and second, both “selfs”, collective and individual, are part of the same structure and interdependent, sharing similar networks and showing a hierarchical organization from a core-self. Both perspectives have some support from current theories in social psychology but are still speculative and faintly supported by empirical evidence. To test this, we designed an experiment using sentences that would activate the individual or collective-self representations in 80 healthy right-handed participants. We use reaction times during a decision-making task, in combination with individualism/collectivism scales and characterize the neural dynamics throughout the experiment using event-related potentials and fronto-parietal informational connectivity networks. Participants reacted slower to the collective than individual self conditions and showed differences in neural activity and information Integration level that distinguished between each type of self. More importantly, the neural integration measure representing the core-self (subtraction between individual and collective wSMI) was only associated with the individualism scores but not collectivism, lending further support for the core-self perspective. We interpret that the collective self, in the broader sense, is a part of the self-concept and therefore probably assembled from the core-self.

## INTRODUCTION

“What is the self?” has been a key question since the beginning of written history, and more recently, for modern social psychology, cognitive neurosciences, and philosophy (Chen et al., 2013). One of the issues that have gained attention in studies of the self is the self-representations of self-concepts (Baumeister, 1999; Gaertner et al., 1999; Kitayama & Park, 2010; Kunda, 1999; Wood et al., 2013). The Self-concept is composed of several components referred to as self-construal that co-exist within an integrated system of self-representations, which in turn, guide behaviour, give meaning, and arrange our experiences (Hardie et al., 2005; E. S. Kashima et al., 2000). At least two fundamental self-representations comprise the self-concept: the collective self and the individual self. The collective self-definition is derived from membership to a social group (Brewer & Gardner, 1996; Y. Kashima et al., 1995; Sedikides & Brewer, 2001); this kind of social identity is associated with attributes and characteristics that come from group membership (Ellemers et al., 2002; Gaertner et al., 2002). The second, the individual self, reflects a person’s self-definition in terms of their unique personal qualities; and it’s a self-definition that is supposed to be independent of group membership (Gaertner et al., 1999, 2002, 2012) and related to the person’s identity (Kunda, 1999; Markus & Kunda, 1986).

Recent work in social psychology and cognitive sciences suggests that the individual self and the collective self are two separate and autonomous self-representations, as has been proposed in the optimal distinctiveness theory (ODT) (Brewer, 2003; Leonardelli et al., 2010). This perspective suggests that both kinds of self are socially and individually adapted to the environment and may have evolved to buffer collective interactions, while the individual differentiate themself from the others. Another approach, however, suggests that our social identity is composed of “many selfs” where the collective self –based on some specific community identity-is only one of them; and that the social identity that holds our identity (individual self), and our self-concepts are heavily influenced by the groups to which they belong (Hogg, 2001). This points to the idea that both types of self-representations aren’t independent or dissociable components and, accordingly, they would be interdependent and possibly hierarchically organized, according to culturally dominant attributes. This has been useful to differentiate between western and eastern societies (Han & Northoff, 2009; Kitayama & Park, 2010). Studies on “working self-concept” suggest that such cognitive disposition corresponds to a sum of attributes that constitute both types of self (Kunda, 1999; Markus & Wurf, 1987). Nevertheless, there’s no direct experimental evidence to support any of these assumptions.

In this same line, but from another perspective, Panksepp & Northoff (2009), points out that “the most basic form of self is, at its core, relational in the sense that it constitutionally allows the selective and adaptive relation of the organisms to their environments” (p.194). On the other hand, when referring to specific neural networks, they refer to the underlying system as the “core-self”. The “proto-Self” (Craig, 2002; A. Damasio & Meyer, 2009; Panksepp, 2005) -the most ancient form of coherent body representation-would be the infrastructure that allows the emergence of a core-Self. Most animals and humans share the first two levels of the self, that is, a proto-self (self-related processing [SRP] as a specific mode of interaction between organism and environment) and core-self (specific neural networks that provide primordial neural coordinates that represent organisms as living creatures) (Panksepp & Northoff, 2009). While the reflective or cognitive self may be reserved for animals that have evolved to infer their cognitive states, at the very least, cetaceans, elephants, birds (e.g., crows), Cephalopoda (e.g., octopuses) higher primates and humans (Birch et al., 2020; Panksepp, 2005).

The reflective self would drive the top-down modulation of the core self between lower and higher aspects of selfhood. In high-order processing, the self-core would mediate between the organismic signals and its physical body image, cognitive, cultural and other environmental contexts, producing a diversity of selfs (personalities), which have been emphasized in philosophical, humanistic and cultural studies, according to Panksepp & Northoff (2009). The core-self may drive and determine the higher forms of self, and if the interaction is deficient, it could generate pathological structures of the personality (Gallagher, 2000; Panksepp & Northoff, 2009).

Complementarily, Feinberg (2000) has proposed that the self is arranged in an organic nested hierarchy. Parts that comprise the lower levels of the hierarchy are physically combined or nested within higher levels to create an increasingly complex set. In the nested hierarchy of a unified mind, lower-order features are incorporated in the mind as “part of,” or nested within a higher-order property. In this way, neural networks from the upper and lower levels of the hierarchy can contribute to sensory awareness and allow for mental unification (an action that is incorporated within the entire hierarchical system through multiple hierarchical levels) (Feinberg & Keenan, 2005). One way to implement these hypotheses in human cognitive neuroscience is through the integration of information via information connectivity measures (King et al., 2013; Koch et al., 2016; Sporns, 2013), allowing to theoretically frame conceptual layers of the self in neural networks information terms and bridge the implementation gap.

Traditionally, from social psychology, self-representations of the self-concept have been studied employing tasks in which participants evoke their kind of self by reading sentences that are related to features linked to each self’s domain. Phrases like “I am a unique and particular person” or “My characteristics are very similar to this group membership”, are used to reflect the individual and collective self, respectively (they are usually answered through a continuous scale with two poles, “yes” or “no”). Previously, each kind of self is experimentally manipulated via distinctive features of the kind of identity. For instance, when the individual self is activated (attribute of “one being creative”), the subject receives negative feedback after having performed a creativity task (“you are little or no creative at all”). A similar way occurs when the collective self is enhanced (“creative ability” of the group to which it belongs): the participant is told that their group is not very or not creative at all. After that, is evaluated, which self-representation is more primary (Gaertner et al., 1999). Other experimental studies about the Self have used images of objects that belong to the participant vs objects belonging to others (Miyakoshi et al., 2007) and the name of its vs other (Gray et al., 2004; Perrin et al., 2005). In these cases, event-related potentials (ERP’s) were associated with self vs no-self conditions (modulation of P300 and N250). In one of the experiments (Chen et al., 2013), participants performed an oddball task with self-relevant stimulus to individual self vs collective self (a circle was used as the target stimulus for each kind of self: each circle contained, on the inside, the subject’s own forename vs subject’s own surname, respectively). Results show that larger P3 amplitudes were evoked by individual self-relevant than collective self-relevant stimuli, while P2 was modulated by both kinds of selfs. However, this research isn’t directly tapping into the cognitive and neural mechanisms that underlie the self-representations but reports on brain correlates associated with each type of self. Another study, carried out by Sul et al., (2012), explored cultural differences in the neural circuits associated with the personal self in contrast with the social self (similar to the individual and collective self). They conducted two experiments exploring the self-reference effect (SRE) paradigm in an fMRI setting, where participants had to memorize words from a list of attributes that described personality traits (personal words) or social identities (social words). They found main effects for targets in areas previously linked with self-processing. Specifically, the left temporoparietal junction was commonly recruited, regardless of the self-aspect among collectivist variables. In contrast, among individualist variables, the left temporoparietal junction was less active when the personal self was considered compared to the social self. These suggest that both self-representations could be part of the same network or structure of the self and that it can be modulated depending on the information that is accessed (individual or social).

Considering all this, the present research aims to identify which cognitive and neurophysiological features from each kind of self, underly such self-representations. We have conceptualized the core self in two alternative accounts; a minimal “subtractive” one, and a fundamental network that drives the other “self’s”. The first account assumes that the core is only a minimal representation that gets revealed if the individual and collective selves are “subtracted” and it is not truly independent but a minimal construct. The second account puts forward the idea of a hierarchy where the core is a driving force to the other selfs. About this second proposal, we hypothesize that core-self hierarchically organizes the different ways of self-representation at several levels, in the way that Feinberg (2000) proposes.

To study the self within a cognitive neuroscience framework, we adapted a specific task from social psychology. We tested whether the self-representations (individual and collective self) are independent dimensions of the self-concept (different networks and process of manipulation of the information), or both correspond to the same cognitive structure, sharing similar networks, being interdependent and hierarchically organized. We expected to find evidence in favour of the second hypothesis. We think both kinds of self are built within a similar architecture, and we predict the collective self to show less neural information as it would reflect information processing for only a portion of the total core-self.

## RESULTS

We used an adapted social psychology paradigm to evaluate our hypothesis; first, we build two types of sentences to activate individual-self and collective-self. Those aimed at activating the individual self were phrases such as “I am hostile”; “I am nice”; “I am spontaneous”, “I am cheerful”, and “I am obstinate”, among others. The sentences to evoke collective-self were similar in their adjectives in terms of length and meaning to individual-self, but were preceded by the phrase “Chileans are” instead of “I am”. The participants had to press a key for yes or no for each stimulus (see Figure 1A). While they responded to sentences, reaction times and EEG activity were measured. EEG neural dynamics recorder was used to obtain ERPs and weighted symbolic mutual information (wSMI, in the way proposed by Canales-Johnson et al., 2020, 2021; Imperatori et al., 2019a; King et al., 2013). We predict that if they are independent of each other, Neural information (wSMI) would be greater for stimuli for the collective self vs. individual self. The underlying idea is that the collective self requires additional information processing as it further compares with information of the individual self before deciding on a statement. On the other hand, if both selfs are built within a similar architecture, we predict the collective self to show less neural information as it would reflect information processing for only a portion of the total core-self.

**Figure 1.**
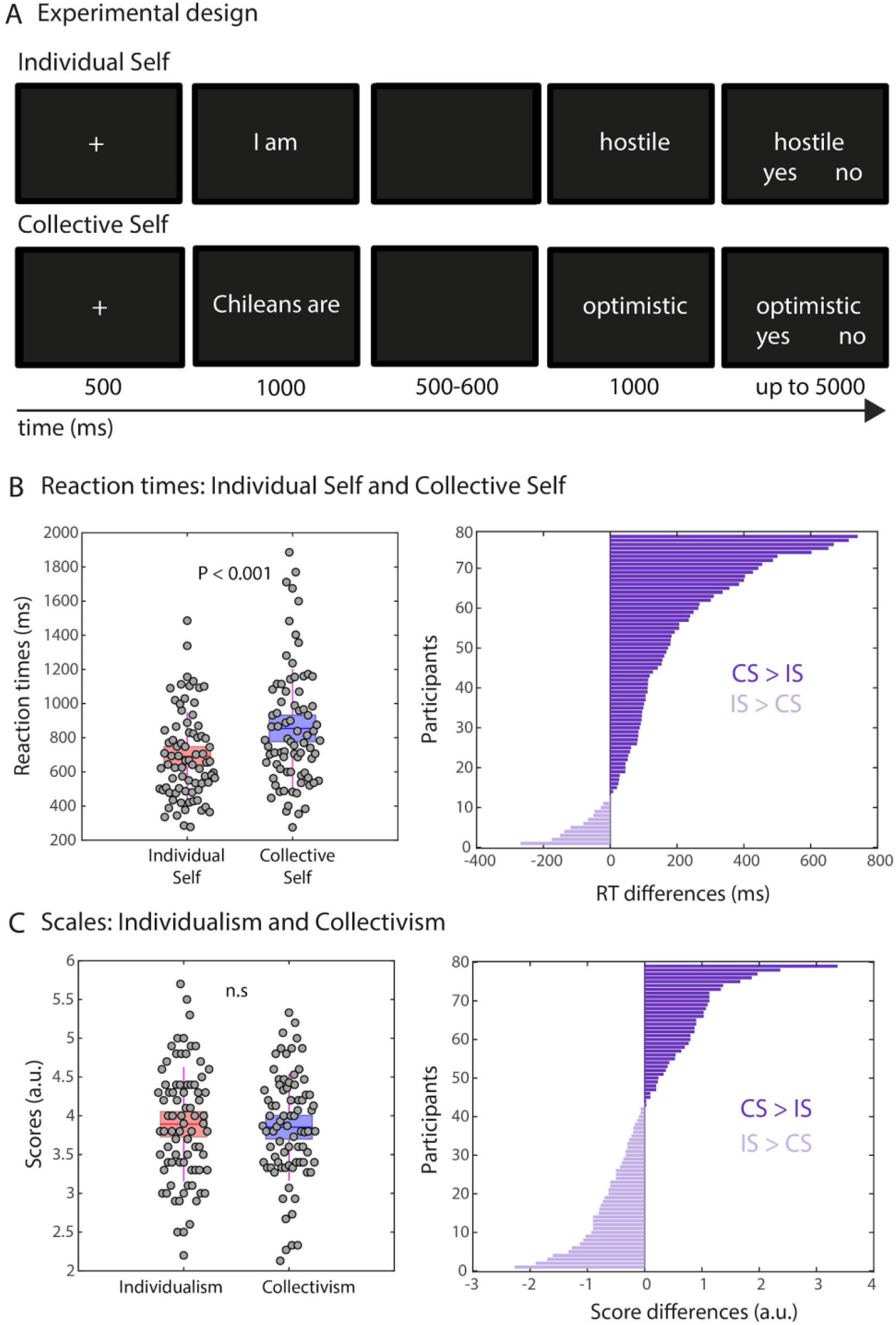
Experimental design and behavioural findings. Individual self-identification was faster than collective self while individualism and collectivism scores were comparable. **(A)** Experimental design. (**B)** Group-level reaction times results (left panel) and individual reaction time differences (right panel). **(C)** Group-level Likert scale results (left panel) and individual score differences (right panel).

### Reaction time (RT) to individual-self and collective-self stimuli

As predicted, the reaction time response to collective-self was longer than individual-self stimuli; *M*(Collective Self) = 857.63ms (*SD* = 345.22) vs *M*(Individual Self) = 695.07ms (*SD* = 258.20), *t*(79) = 7.10, *p* < .001, Cohen’s *d* = 0.53. Since faster reaction times are associated with automaticity, this would be an indicator of greater accessibility to our cognitive schemes and categories related to such events (Bargh, 1989; Bargh & Williams, 2006; Higgins, 1996; Shiffrin & Schneider, 1977). Secondly, our social comparison is egocentric and therefore requires Self-judging first before others (Dunning, D., & Cohen, G. L. 1992; Dunning, D., & Hayes, A. F. 1996).

### Individualism and Collectivism Scale

To have a behavioural measure that could be associated with individual-self and collective-self, we apply the Auckland Individualism and Collectivism Scale (AICS; Shulruf, et al., 2007). Two measures were obtained: individualism scale (α = 0.76) from 10 items that showed a corrected item-total correlation measure > 0.25 (3 items were eliminated for being below this threshold); and collectivism scale from 10 items (α = 0.81), from a total of 15 items (all shown a corrected item-total correlation measure > 0.27; the rest were eliminated). No reliable differences were observed between both measures, *t*(78) = 0.346, *p* = 0.73, thus, as a group there was a wide range of differences between Individualist and collectivist traits; *M* = 3.89 (*SD* = 0.74) and *M* = 3.85 (*SD* = 0.69) but no imbalance.

### ERP of individual-self and collective-self representations

We first report a modulation of P2 and P3 event-related potentials (ERP) to individual and collective attributes of the self, (in agreement with Chen et al., 2013; Gray et al., 2004; Miyakoshi et al., 2007; Perrin et al., 2005). We performed a cluster-based permutation test in two predefined windows of interest: an early window from 0 to 280 ms; and a late time window from 300 to 600 ms. In the early window, we observed a reliable anterior electrode cluster between 180-280 ms after stimuli presentation, showing higher activity for the individual than collective self condition (P < 0.001; Figure 2A). Importantly, these differences were highly consistent at the single-participant level (Figure 2B). Finally, source reconstruction analysis showed that the scalp-level voltage differences were primarily driven by bilateral frontal and temporal-left brain regions (P < 0.001; Figure 2C). Similarly, in the late time window, we observed an increase in voltage in the individual-self condition as compared to the collective self-condition between 350-450 ms (Figure 2D). Again, these differences were highly consistent across participants (Figure 2E), and source reconstruction analysis showed that the scalp topographies were mainly driven by frontal bilateral and temporal-left sources (Figure 2F). Overall, these results confirm that early (P2) and late (P3) ERP neural signatures consistently distinguish individual and collective attributes of the self. To test if P2 and P3 are associated with the individual and collective self (Chen et al., 2013) we tested an association with the scales to probe the psychological and cultural traits of individualism and collectivism. We found no reliable correlations (n=80) for individual self P2 (180-280ms) and P3 (350ms - 450ms) with Individualism Scale, Pearson’s *r* = -0.10, *p* = 0.37 and Pearson’s *r* = 0.02, *p* = 0.86; to collective self P2 (180-280ms) and P3 (350ms - 450ms) with Collectivism Scale, Pearson’s *r* = 0.01, *p* = 0.90 and Pearson’s *r* = 0.18, *p* = 0.11.

**Figure 2.**
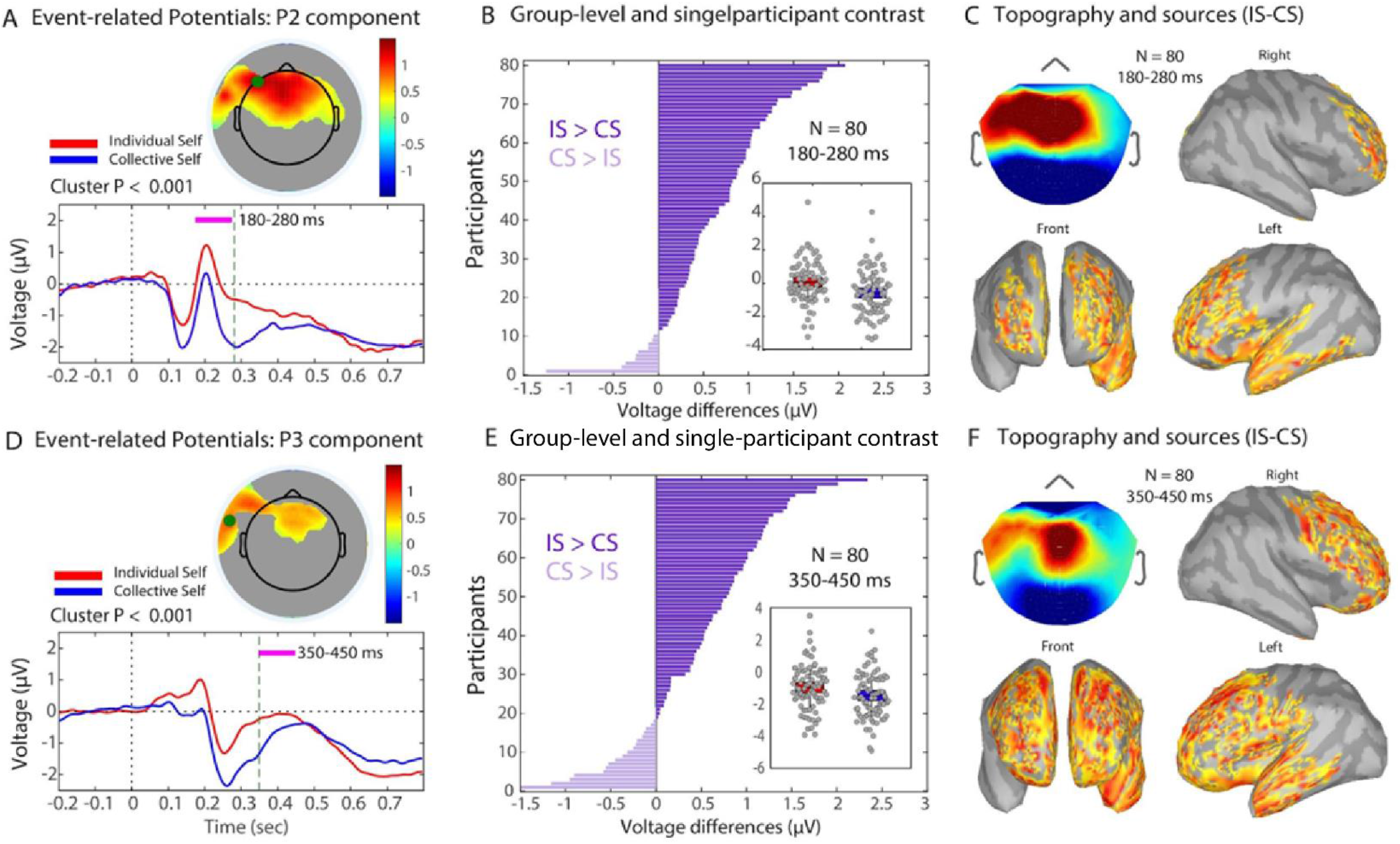
ERP results (A). Early (180-200ms) and late (350-450ms) neural activity components showed higher brain activity for the individual than collective Self. *Left panel:* Evoked-related potentials of Individual Self (IS; red) and Collective Self (CS; blue) highlighting the P2 component window (180-280 ms; grey rectangle. *Right panel:* Topographical differences between Individual and Collective Self (IC-CS). (C) Group-level statistics of P2 amplitudes of IS (red) and CS (blue). The horizontal line inside the box represents the mean; the box represents the standard error of the mean (S.E.M) and the vertical line (magenta) represents the standard deviation (S.D). **(C)** P2 voltage differences (IS - CS) per participant. (D-E-F) Same as (A-B-C) but for the P3 component (350-450 ms).

### Frontoparietal information integration and self representations

The fact that representing different features of the self involves multiple neurocognitive sub-processes such as embodiment, self-reference, metacognition, and autobiographical memory (Botzung et al., 2008; Chiao et al., 2009; A. R. Damasio et al., 2000; Ellis-Hill & Horn, 2000), suggests that individual and collective self-representations might require the integration of information across multiple brain networks. Then, we reasoned that the neural underpinnings of the self may be better characterised by changes in connectivity in a distributed network of brain regions. We hypothesized that a neural metric specifically indexing long-distance neural information integration (wSMI; Canales-Johnson et al., 2020, 2021b; Imperatori et al., 2019a, 2019b; King et al., 2013) could in principle capture modulations in information dynamics between the two types of self. Although studies showing connectivity modulations of self-related phenomena are scarce, there is some preliminary evidence suggesting that changes in frontoparietal alpha- and beta-band connectivity are associated with phenomenological features of the self such as the sense of agency (Alzueta et al., 2020; Kang et al., 2015).

Following this preliminary evidence, we contrasted the dynamics of information integration in the beta and alpha bands across frontal and parietal electrodes between individual and collective self. wSMI differences were computed in two-time windows of interest: an early window (180 to 280 ms) and a late window (350 at 450ms), matching the time windows used for P2 and P3 ERP analyses, respectively (Figure 2). In the frontoparietal beta-band wSMI, a RANOVA between self representation (individual vs collective) and time window (early vs late) revealed a reliable main effect for self representation (*F*_1,79_ = 781.09, *P* < 0.001, Cohen’s *d* = 6.28) with higher wSMI for the individual self, and a reliable main effect of the time window (*F*_1,79_ = 6.39, *P* < 0.05, Cohen’s *d* = 0.58) with higher wSMI for the early window. No reliable interaction was observed between windows (*F*_1,79_ = 0.10, *p* = 0.75).

In contrast to beta-band wSMI results, RANOVA in the frontoparietal alpha-band wSMI (self representation x time window) revealed no differences between individual and collective self (*F*_1,79_ = 3.11, *ns*), but only a main effect of the time window (*F*_1,79_ = 347.31, *P* < 0.001, Cohen’s *d* = 3.79) with higher wSMI for the early window. Again, no reliable interaction was found (F = 0.049, *p* = 0.83). These results suggest that the information contained in fast frequencies (beta more than alpha) across distributed networks might be relevant for indexing different representations of the self.

Finally, we explored the possible dependencies between each type of self to evaluate if they could belong to a partially shared common network. For this, a bivariate correlation was performed between the wSMI individual self (τ = 6 ms) and wSMI collective self (τ = 6 ms) an early window for both. We find a strong association, Pearson’s *r*= 0.86 (N=80), p < 0.001, Cohen’s d = 3.37. From these results, we can conclude that individual and collective self can be neurally marked in early windows beta-band frontoparietal networks with wSMI.

### Core-self, information integration and individualism/collectivism scales

Not enough evidence for associations was found between beta-band wSMI (early window) measures – individual and collective self *–* and self-representations scales (Individualism and Collectivism): to individual wSMI with Individualism scale, Pearson’s *r*= 0.06 (N=80), *p* = 0.59, and collective scale, Pearson’s *r*= 0.008 (N=80), *p* = 0.95; to collective wSMI with Individualism scale, Pearson’s *r*= - 0.097 (N=80), *p* = 0.40, and collective scale, Pearson’s *r*= 0.044 (N=80), *p* = 0.70. Since the current findings -that individual self-concept showed higher information sharing on beta-band than collective-self and that wSMI to collective and individual self are strongly correlated, we can suggest that different representations of the self may be part of one same common factor. We propose a core-self that contains all our self-representations in the same network. Each of them is activated when stimulated by the environment. If our conception is right, when subtracting the collective self from the individual self, we should observe information that contains a more robust link with the core-self. Since more wSMI information is associated with an individual self, then we suggest that the remainder of the subtraction is compatible with the idea of an individual core-self. This proposition is consistent with current reports of individual self primacy in Western cultures (Han, 2022). To corroborate this assumption, we performed a linear regression model using the individualism and collectivism scale, predicting the wSMI beta-band of core-self as predictors, for both early and late windows. The effect was found in an early window on a simple linear regression, *F*(2, 76)= 4.32, *p* < 0.05, *R*^*2*^= 0.10., and uniquely for individualism scale, β = 0.31, *t* = 2.79, *p* < .01, Cohen’s *d* = 0.65; but not for collectivism scale, β = 0.09, *t* = 2.9, *p* = 0.39, *ns*. For the late window, there was no reliable evidence, ANOVA, *F*(2, 76)= 0.12, *p* = 0.89, *ns*., Figure 4 shows the neural marker and the individualism and collectivism scale in Pearson correlations as a direct visual representation of the association between factors.

**Figure 4.**
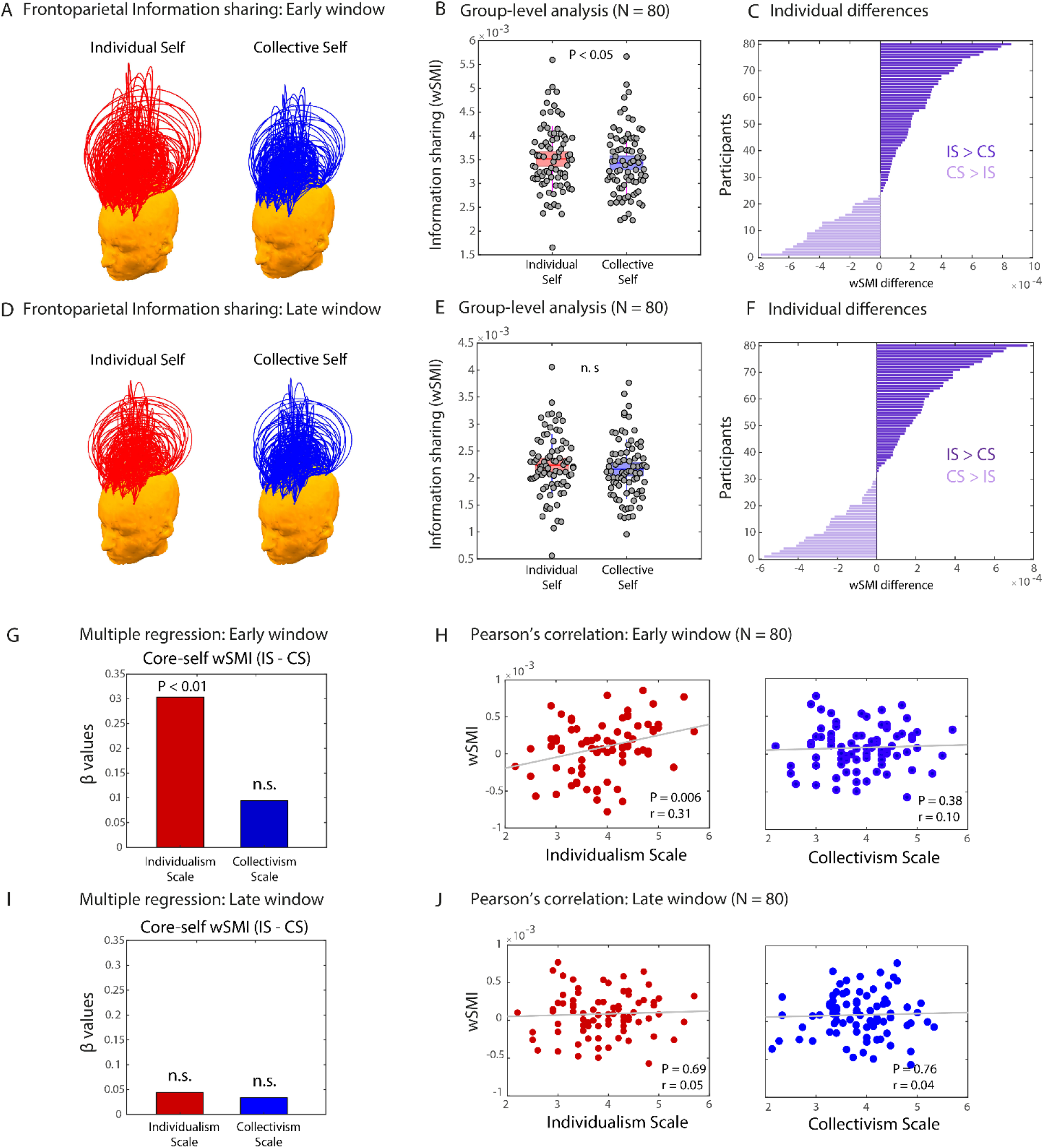
Core-self frontoparietal information-sharing and individual and collective scales. Higher frontoparietal information sharing for individual-self than collective-self in the early but not late window; reliable association between the individualism scale and frontoparietal information sharing but not Collectivism Scale, only in the early window. Frontoparietal information sharing (wSMI) in the beta band in the individual self and collective self conditions. **(A, D)** Each arc represents a functional connection between a pair of electrodes, the height of the arc represents the value of the wSMI or the individual (red) and collective (blue) self for the **(A)** early window (180-280 ms) and **(D)** late window (350-450 ms). Beta-band wSMI calculated between each frontal and parietal electrode. wSMI values within frontal and parietal electrodes regions not included to only capture information integration between distant pairs. **(B, E)** Group-level differences in beta-band wSMI between individual and collective self conditions in the **(B)** early and **(E)** late window. **(C, F)** Individual differences in beta-band wSMI for each participant in the **(C)** early and **(F)** late window. **(G)** Beta coefficients of multiple regressions using Individualism Scale and Collectivism Scale as predicted variable and core-self wSMI as regressor during Early Window. **(H)** The direct association is represented by Pearson’s correlation during Early-window between core-self wSMI and Individualism Scale (left panel) and Collectivism Scale (right panel). **(I)** Beta coefficients of multiple regressions using Individualism Scale and Collectivism Scale as predicted variable and core-self wSMI as regressor during Late Window. **(J)** The direct association is represented by Pearson’s correlation during Late-window between core-self wSMI and Individualism Scale (left panel) and Collectivism Scale (right panel).

## Discussion

In this study, we aimed to use cognitive and neurophysiological tools to differentiate between possible competing hypotheses about the cognitive structure of the self in its collective and individual representations. We asked whether the self is represented by separate networks or is hierarchically organized. For this, we adapted a typical social psychology task to “activate” both types of self-representations and experimentally confirmed expected effects about the collective and individual “selfs”, that when participants accessed individual self-schema through the context linked to individual identity, the reaction times were faster than when they responded to the same stimulus but within the context associated with the collective self. This is consistent with the known effect of accessibility on our information and that is strongly compatible with cognitive schemes of the self (Bargh, 1989; Bargh & Williams, 2006; Higgins, 1996; Shiffrin & Schneider, 1977). Second, we showed, via ERP analysis, that the P2 and P3 components were higher in activity for individual-self than collective-self representations (Chen et al., 2013; Gray et al., 2004; Miyakoshi et al., 2007; Perrin et al., 2005). Thirdly, we show a specific association between the individualism scale with neural information integration measures in the early processing time window, but not in the late window and no association with the collectivism scale. The individualism scale positively associated with the core-self built the evidence in favour of the “selfs” being part of the same cognitive structure, sharing similar neural networks, and being interdependent and hierarchically organized (Feinberg, 2000; Feinberg & Keenan, 2005). the Collective self showed less neural information than the individual Self, and when subtracted, the individualism scale explained the neural variance of the core Self while the collectivism scale did not. We show the core self as a unifying construct for the representation of each self under a precise hypothesis (Panksepp & Northoff, 2009).

The specific neural network that constitutes the minimal neural representation allowing an organism to relate to its environment is referred to as ‘core-self’. We use here a neural metric relying on information connectivity that is better framed theoretically from a point of information processing systems. The approach takes a conceptual knife to differentiate putative distinct processes at play when hearing stimuli related to the individual self or the collective self, assuming either a separate or nested information-sharing network. Furthermore, by taking advantage of the behavioural scales, the participant can metal represent their identity when forced to allocate themselves to a specific category, and when we probe the neural information from the cognitive process to the scales’ outcome we link back the neural echo of the processes to the reported self representation in the questionnaire. This allowed us to use the neural information to differentiate between alternative hypotheses, lending support to the idea of the core-self as the remaining neural network after the collective-individual subtraction.

Further evidence towards the need to frame neural measures to possible underlying information of the networks supporting the self neural representation is specifically supported by the differences in the association between the specific individualistic and collectivistic scales and the neural activity (ERPs) or neural information (wSMI) measures. Simply looking for a neural correlate such as brain activity in EEG -ERPs- to associate with the Scales revealed no association and had no theoretical motivation, despite reflecting a neural difference between the response to the individualistic and collective self representation (Figure 2). In contrast, the neural information measure of the network captured a specific dissociation of individualistic and collectivist Scales, allowing us to interpret it as evidence of a specific organization of the self-core concerning the individual and collective selfs. Thus, we believe that the output of something complex such as the self, which arguably integrates multiple neurocognitive processes, is better reflected by a connectivity metric of long-distance neural information integration such as wSMI, and not by simple activation metrics derived from averaged voltage activity such as the classic ERPs (Canales-Johnson et al., 2020).

Several methodological arguments strengthen the results and interpretation and their ecological validity. First, previous studies have oversimplified the stimuli delivery without validating or defining individualism or collectivism by contextual information, as we do in the present study. A second aspect of the bonafide design is the testing of ∼80 participants and the calculation of the direction and strength of effects in each participant, allowing for a decent evaluation of the degree of association between variables (Schönbrodt & Perugini, 2013) and lending a moderate level of trust in the association between behaviour and neural signals (Hulley et al., 2001). And thirdly, we combined the task behavioural and neural output per participant with their scales scores and specific contrastive statistical design to test the hypotheses at hand.

Alternative interpretations of the results point to two interesting conceptual differentiations, one is the primacy of the individual self, that in our results shows an association between the neural echo of the core self to the individualism scale but not the collectivism scale, suggesting a core representation of the self in individualistic terms both at the processes and possibly at the identity levels (cultural primacy of the self). An alternative interpretation could be that regardless of the cultural context, collectivistic or individualistic, the neural information captures the core as an essential representation, where other representations may rely on (layered “selfs”). It could be that the core-self ties and dependencies with the individual and collective selfs are culturally influenced (different in more eastern collectivist societies from western more individualistic societies), and hence the neural underpinnings of the core-self with the individualism scale found here are reversed in more collectivist samples. The core-self may be the true reflection of the cultural context, built throughout development and societal constraints.

These findings, however, lead us back to the discussion of whether the core-self is constituted by a strong primacy of individual aspects of the self (our particular traits, personal characteristics, our individual identity), or rather, reflects the contextual predominance of a specific type of moulding of the self from the dominant identity traits of the culture in which it develops. If the self-core is a kind of active container that integrates memories, experiences and affective states of the relationship between the self and the world, perhaps then it is what Cooley (1902) called the “looking-glass self”. That is, the way our culture and interaction with others, as if they were a mirror, influences the construction of our own identity, thus reflecting our rich social life. Given the above, one way to experimentally approach the question of whether the conformation of our identity, and accentuated features of our self-core, would be to test if there are essential aspects of individuality or selfhood, or if it is more plastic and dependent on the influence of the dominant context. Perhaps, testing these ideas in highly collectivistic societies would act as a counterpoint to the understanding of the “selfs”, reasoning that if the predominant traits of self-core are collectivistic rather than individualistic (the latter, found in this study), we could interpret that the self-core is the result of the primacy of its context.

A long tradition in Social Psychology has investigated how we represent ourselves and how we approach the study of the self. Through experiments and social cognition theory, it has been proposed that the self emerges when people’s self-concept (for example, individual or collective) is threatened or psychologically amplified. From this perspective, the “self” may be construed as an organized collection of feelings and beliefs about oneself. It influences the way we process, utilize and internalize information about the world in light of motivations, emotional states, self-evaluations and self-schemas (Baumeister, 1999; Kruglanski & Higgins, 2003). On the other hand, the concept of self provides an essential point of contact between theories of personality and theories of social behaviour (Brewer, 1991), and in this sense, the self is not a one-dimensional construct but rather entails various extensions, and each of us has many concepts of himself (Kunda, 1999; Markus & Kunda, 1986). Although research in recent decades has provided some plausible frameworks that allow us to understand how we represent ourselves, until now, there is scarce theorizing consistent with neurocognitive mechanisms and/or based on experimental evidence, and by far little or nothing integrated into social cognition. We believe that a bridge to integrating the theory and experimental evidence from social psychology with theoretical and empirical models proposed by neuroscience and the philosophy of mind may need to be built before further experimental work. The present study, meanwhile, has specifically addressed the question of selfhood, its self-representation of self-concepts, and possible neurocognitive representations that underlie its integration into social life, strengthening the concept of core-self for further theorizing and subsequent experimentation.

## METHODS

### Participants

Eighty healthy human participants (40 female) aged 18 to 26 (*M*=20.5, *SD*=2.09), all students of Psychology, recruited from the University, participated in this experiment for monetary compensation. All participants had a normal or corrected-to-normal vision and had no history of head injury or physical and mental illness. This study was approved by the local ethics committee of the Universidad Diego Portales, Santiago de Chile, and written informed consent was obtained from all participants after an explanation of the experimental protocol.

### Experimental task

Participants performed an adapted version of the motivational primacy of the self paradigm (Gaertner et al., 1999) from Social Psychology. The task was that each participant had to answer “yes” or “no” to 200 sentences that alluded to the individual self-concept (e.g., “I am intelligent”) and another 200 sentences to the collective self-concept (“Chileans are united”). These phrases appear on a screen 500 ms after a fixation cross. Later, the options “yes” or “no” associated with a key appeared, which had to be selected.

### Procedure

Statements were constructed with qualifying adjectives to activate each type of self, based on individual or collective self-representation. A total of 272 sentences were constructed to evoke the individual self, such as, “I am generous”; “I am proud”; “I’m a liar”; “I am ironic”; among others. Likewise, phrases were generated to trigger the collective self (e.g., “Chileans are friendly”; “Chileans are liars”; “Chileans are fun”; “Chileans are resentful”). These stimuli were validated in a sample of 411 students (similar in characteristics to subsequent study participants), based on representativeness as positive and negative valence (with Likert scales from 1 to 7). To choose the final stimuli for the experiment, the t-test scores for a sample were standardized. The μ value used was 4, which corresponds to neutral evaluation. Scores higher than 2 and small values of -2 were considered large values, that is, values that are at least 2 standard errors away from the zero points (value 4 on the original scale). With this criterion, 200 stimuli were obtained for each self.

### EEG recording and processing

EEG was recorded and digitized with a GES300 Electrical Geodesic amplifier at 500 Hz and 129-channel saline-based wet HydroCel Geodesic sensor nets. During recording, the signal was pass-band filtered in the 0.01-100 Hz range. During the acquisition electrode, Cz was used as a reference and impedances were kept under 50 kΩ. The EEGLAB (Delorme and Makeig, 2004) and ERPLAB (Lopez-Calderon and Luck, 2014) Matlab toolboxes were employed for offline data processing. The signal from before and after the task, and between blocks was removed and the remainder was filtered between 0.5-30 Hz. An independent component analysis (ICA) was performed on the continuous EEG signal. Components attributed by visual inspection to eye blinks, eye movements, heartbeats, and line noise were removed. The clean signal was average referenced and epoched from 200 ms before to 800 ms after the adjective presentation. On the 91 centermost electrodes, trials containing voltage fluctuations exceeding ± 100 μV or sample-to-sample differences exceeding ±30 μV were rejected. Trials were averaged by collectiveness and valence of stimuli.

### ERP analyses

Trials were compared in pairs of experimental conditions using spatiotemporal clustering analysis implemented in FieldTrip (Oostenveld et al., 2011) using dependent sample t-tests. Although this step was parametric, FieldTrip used a nonparametric clustering method to address the multiple comparisons problem. *t* values of adjacent spatiotemporal points with p<0.05 were clustered together by summating them, and the largest such cluster was retained. A minimum of two neighbouring electrodes had to pass this threshold to form a cluster, with neighbourhood defined as other electrodes within a 4 cm radius. This whole procedure, i.e., calculation of t values at each spatiotemporal point followed by clustering of adjacent t values, was repeated 1000 times, with recombination and randomized resampling before each repetition. This Monte Carlo method generated a non-parametric estimate of the p-value representing the statistical significance of the originally identified cluster. The cluster-level t value was calculated as the sum of the individual t values at the points within the cluster. Spatiotemporal clustering was always carried out within a restricted time window of -200 to 800 ms after stimuli presentation.

### EEG source reconstruction

To visualize the brain origins of ERPs (Figure 3), cortical sources of subject-wise averaged ERPs were reconstructed using Brainstorm (Tadel et al., 2011). The forward model was calculated using the OpenMEEG Boundary Element Method (Gramfort et al., 2010) on the cortical surface of a template MNI brain (colin27) with 1 mm resolution. The inverse model was constrained using weighted minimum-norm estimation (Baillet et al., 2001) to calculate source activation. To plot cortical maps, grand-averaged activation values were baseline corrected by z-scoring the baseline period (−200 to 0 ms window) to each time point, and spatially smoothed with a 5 mm kernel. This procedure was applied separately for the individual-self, and collective-self conditions, and averaged across the significant time windows shown in Figure 3A and D.

### Information sharing analysis: weighted symbolic mutual information (wSMI)

To quantify the information sharing between electrodes we computed the weighted symbolic mutual information (wSMI) (King et al., 2013; Imperatori et al., 2019). It assesses the extent to which the two signals present joint non-random fluctuations, suggesting that they share information. wSMI has three main advantages: (i) it allows for a rapid and robust estimation of the signals’ entropies; (ii) it provides an efficient way to detect non-linear coupling; and (iii) it discards the spurious correlations between signals arising from common sources, favouring non-trivial pairs of symbols. For each trial, wSMI is calculated between each pair of electrodes after the transformation of the EEG signals into sequences of discrete symbols defined by the ordering of k time samples separated by a temporal separation tau. The symbolic transformation depends on a fixed symbol size (k = 3, that is, 3 samples represent a symbol) and a variable tau between samples (temporal distance between samples) which determines the frequency range in which wSMI is estimated. The frequency specificity f of wSMI is related to k and tau as:

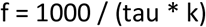

As per the above formula, with a kernel size k of 3, tau values of 12 ms hence produced a sensitivity to frequencies below 30 Hz with and spanning the beta-band (∼15-28 Hz).

wSMI was estimated for each pair of transformed EEG signals by calculating the joint probability of each pair of symbols. The joint probability matrix was multiplied by binary weights to reduce spurious correlations between signals. The weights were set to zero for pairs of identical symbols, which could be elicited by a unique common source, and for opposite symbols, which could reflect the two sides of a single electric dipole. wSMI is calculated using the following formula: where x and y are all symbols present in signals X and Y respectively, w(x,y) is the weight matrix and p(x,y) is the joint probability of co-occurrence of symbol x in signal X and symbol y in signal Y. Finally, p(x) and p(y) are the probabilities of those symbols in each signal and K! is the number of symbols - used to normalize the mutual information (MI) by the signal’s maximal entropy. The time window in which wSMI was calculated was determined based on the significant time window observed in the ERP contrast of Figure 2: early window (825-1675 ms) and late window.

## Statistics

Statistical analyses were performed using MATLAB (2016a), Jamovi (version 0.8.1.6; open source; https://www.jamovi.org), and JASP (version 0.8.4) statistical software.

